# Placental epigenetics for evaluation of fetal congenital heart defects: Ventricular septal defect (VSD)

**DOI:** 10.1101/355768

**Authors:** Ray Bahado-Singh, Samet Albayrak, Rita Zafra, Alosh Baraa, Avinash M Veerappa, Deepthi Mahishi, Nazia Sayed, Nitish K Mishra, Chittibabu Guda, Rouba Ali-Fehmi, Uppala Radhakrishna

**Affiliations:** Department of Obstetrics and Gynecology, Oakland University William Beaumont School of Medicine, Royal Oak, MI, USA.; Department of Obstetrics and Gynaecology, Wayne State University School of Medicine, Detroit, MI, USA; Department of Obstetrics and Gynaecology, Icahn School of Medicine at Mount Sinai, New York, NY, USA; Department of Pathology, Wayne State University School of Medicine, Detroit, MI, USA; Department of Studies in Genetics and Genomics, Laboratory of Genomic Sciences, University of Mysore, India; Biotechnology, Nirma Institute of Science, Nirma University, Ahmedabad, India; Department of Genetics, Cell Biology & Anatomy, College of Medicine, University of Nebraska Medical Centre Omaha, NE, USA

**Keywords:** Epigenetics, DNA methylation, biomarkers, Ventricular septal defect, placental biomarkers, Genome-wide association, Genomics

## Abstract

Ventricular Septal Defect (VSD), the most common congenital heart defect, is characterized by a hole in the septum between the right and left ventricles. The pathogenesis of VSD is unknown in most clinical cases. There is a paucity of data relevant to epigenetic changes in VSD. The placenta is a fetal tissue and is a potentially useful surrogate for the evaluation of fetal organ development. To understand epigenetic mechanisms that may play a role in the development of VSD, a genome-wide DNA methylation assay of the placentas of 8 term subjects with isolated VSD and no known or suspected genetic syndromes and 10 normal controls was performed using the Illumina HumanMethylation450 BeadChip assay. The study identified a total of 80 highly accurate potential epigenomic markers in 80 genes for the detection of VSD; area under the receiver operating characteristic curve (AUC ROC) = 1.0 with significant 95% CI (FDR) p-values < 0.05. The biological processes and functions for these differentially methylated genes are known to be associated with heart development or heart disease, including cardiac ventricle development (*HEY2, ISL1*), heart looping (SRF), cardiac muscle cell differentiation (*ACTC1, HEY2*), cardiac septum development (ISL1), heart morphogenesis (*SRF, HEY2, ISL1, HEYL*), Notch signaling pathway (*HEY2, HEYL*), cardiac chamber development (*ISL1*), and cardiac muscle tissue development (*ACTC1, ISL1*). The study also identified eight microRNA genes that have the potential to be biomarkers for the early detection of VSD including miR-191, miR-548F1, miR-148A, miR-423, miR-92B, miR-611, miR-2110, and miR-548H4. To our knowledge this is the first report in which placental analysis has been used for determining the pathogenesis of and predicting CHD.

## Introduction

Congenital Heart Defect (CHD), affects nearly 40,000 births per year in the United States [1, 2]. Ventricular Septal Defect (VSD) is the most common congenital heart disease (CHD) and occurs in approximately 1 in 500 live births [3–5]. The frequency is even more common in prenatal life. The common risk factors for VSD include family history, ethnicity, and a few genetic disorders. Data indicate that genetic mutations of cardiac developmental genes such as *NKX2-5*, *SMAD3*, *NTRK3*, *GATA6*, *TBX2*, *TBX18*, *ATA6*, and *TBX2*, unique copy number variations [6], and chromosomal aneuploidy such as trisomy 13, 18, or 21 can play an important role in the etiology [7] of VSD. Targeted disruption of the CHF1/Hey2 locus in mice resulting in VSD has also reported [8]. Despite these known genetic disruptions, in the clinical setting the cause of VSD remains unknown in the great majority of cases.

DNA methylation is one of many epigenetic mechanisms and is the most widely studied. Aberrant DNA methylation is a significant player in the transcriptional regulation of genes and has been implicated in many complex and common diseases including cancer, diabetes, or psychiatric disorders. While insufficiently well studied, emerging evidence suggests that DNA methylation may have an important role in both normal and pathologic heart development [9]. Methylation of CpG islands is thought to have its biological effect by displacing transcription binding factors, and instead attracting methyl binding factors that condense DNA and suppress gene expression. Environmental factors such as maternal diet, smoking and alcohol exposure profoundly affect DNA methylation. These factors are known to be risk modifiers for CHD development [1, 10, 11]. Further, signal DNA methylation changes have been demonstrated in the cardiac tissue DNA of CHD fetuses [12].

While methylation is known to be tissue specific it is not tissue exclusive. Several recent studies have shown correlation between CpG methylation specific cytosine loci across different tissues [13, 14]. This has raised the prospect for example of using the DNA methylation status in easily accessible tissues such as blood leucocytes to evaluate methylation status and by inference gene activity and disease status, in inaccessible organs such as the brain [15]. Epigenetic markers in the blood leucocytes have thus been used to detect psychiatric disorders such as schizophrenia [16]. The degree of correlation in cytosine methylation across different tissues is thought to vary based on the specific tissues involved.

Currently, there are no proven biomarkers available in clinical practice for the pre-or post-natal detection of congenital heart defects (CHDs). Given the clinical significance of CHD and the frequency of missed or late diagnosis, this is a major deficiency. In our prior pilot data [17, 18] cytosine methylation status of blood leucocytes was found to be a potentially useful molecular biomarker for the detection of multiple different categories of CHD. This is consistent with the previously referenced attempts to use epigenetic signatures in leucocytes to detect psychiatric disorders [16]. The placenta, like newborn blood that was used in our prior study is a fetal tissue. We therefore reasoned that although not identical, there was likely to be a significant minority of cytosine loci in which parallel epigenetic modifications could be demonstrated in cardiac development genes in general and genes related to ventricular development in the placenta from VSD pregnancies. We had previously demonstrated this phenomenon in multiple categories of CHD using blood (leucocyte) from affected newborns as the surrogate tissue for detecting CHD [17, 18]. Our objective therefore in this proof of concept study was to evaluate the utility of cytosine methylation in placental DNA to elucidate the pathogenesis of and for the detection of isolated non-syndromic VSD. Further we used cytosine methylation to investigate the molecular pathogenesis of non-syndromic VSD based on the associated genes and gene pathways that were differentially methylated in the VSD cases.

MicroRNA (miRNA) is another important epigenetic mechanism and exerts control over DNA methylation and suppresses gene expression among other functions. Recent data suggest and important role for miRNA in CHD development [19]. We therefore also evaluated methylation status of known microRNA genes in-liu of measuring actual miRNA levels. Given that DNA methylation status is known to correlate with gene expression, this approach can be used to identify miRNAs that are involved in cardiac development and thus further elucidate the mechanism of VSD pathogenesis. To our knowledge, there are no prior reports using placental molecular analysis for the detection of investigation of CHD pathogenesis.

## Materials and methods

Tissue samples for analysis: Genomic DNA was isolated from archived formalin-fixed, paraffin-embedded tissue blocks (FFPE) using Puregene DNA Purification kits (Gentra systems^®^ MN, USA) according to manufacturer’s protocol. The tissue samples were taken from the placenta of VSD subjects immediately after birth. DNA from archived paraffin-embedded tissue is a suitable template and has been used previously for genome-wide DNA methylation profiles using the Infinium HumanMethylation450 BeadChip assay. Importantly, it has been demonstrated that it is possible to achieve high-performance outcomes using FFPE-derived DNA in genome mapping arrays [20]. FFPE derived DNA captures on average more than 99% of the CpG sites on the array. The institutional review board of the Wayne State University School of Medicine approved the study.

### Genome-wide methylation analysis using the HumanMethylation450

The HumanMethylation450 (Illumina, Inc., California, USA) contains >485,000 CpGs per sample in enhancer regions, gene bodies, promoters and CpG islands at a singlenucleotide resolution and requires only 500 ng of genomic DNA. Many studies have previously used Illumina Infinium technology to assess DNA Methylation changes in the placenta associated within fetal disorders [21]. DNA was bisulfite converted using the EZ DNA Methylation-Direct Kit (Zymo Research, Orange, CA), and fluorescently-stained BeadChips imaged by the Illumina iScan. Data were analyzed with Illumina’s Genome Studio methylation analysis package program. The methodology has been previously detailed earlier [17].

As previously reported, to avoid potential confounding factors, probes associated with sex chromosomes and/or containing SNPs in the probe sequence (listing dbSNP entries near or within the probe sequence, i.e., within 10 bp of the CpG site) were excluded from further analysis [22–24]. Probes targeting CpG loci associated with SNPs near or within the probe sequence may influence corresponding methylated probes [25]. The remaining CpG sites were analyzed.

Statistical and Bioinformatic analysis. Differential methylation was assessed by comparing the ß-values per individual nucleotide at each CpG site between VSD subjects and controls. The p-value for methylation differences between case and normal groups at each locus was calculated [26]. Filtering criteria for p-values was set at <0.05 and <0.01 to identify the most discriminating cytosine’s or the most important differentially methylated regions of the genome. P-values are calculated with False Discovery Rate (FDR) correction for multiple testing (<u>Benjamini-Hochberg test</u>). Further analysis of the differentially methylated genes was conducted for potential biological significance. Receiver Operating Characteristic (ROC) and Area Under Curve (AUC) with R was calculated to determine the diagnostic accuracy of specific cytosine loci differentiating CHD from control groups. Data are normalized using the Controls Normalization Method.

The most significantly differentially methylated CpG sites were selected based on pre-set cutoff criteria of ≥2.0-fold increase and/or ≥2.0-fold decrease with Benjamini - Hochberg False Discovery Rate (FDR) p<0.01. Multiple CpG sites within a gene were resolved by selection of the CpG with the highest fold-change ranking and the lowest p-value. Fold changes in methylation variation were obtained by dividing the mean ß-value for the probes in each CpG site by that of the normal controls. After this filtering, a threshold was set to select ROC curves based on sensitivity plotted against specificity using different ß-value threshold at each CPG locus for VSD prediction were constructed. The corresponding AUC (95% CI) was calculated to select disease marker at each CpG locus and corresponding gene. In the case of multiple differentially methylated cytosine loci in the same gene we use the locus with the highest discriminating power as defined. To avoid potential experimental confounding, various statistical modeling was used. Markers with AUC ≥0.80 and significant 95% CI and FDR p-value < 0.005 were further used to generate heatmap and pathway analysis.

Gene ontology analysis and functional enrichment. Pathway analysis was carried out using Ingenuity Pathway Analysis (Ingenuity Systems, www.ingenuity.com) using differentially methylated genes (at FDR p-value < 0.01). Literature data mining for co-occurrence of gene names and keywords of interest was performed using Chilibot (www.chilibot.net). Only genes for which Entrez identifiers were available were used in the Pathway analysis. Over-represented canonical pathways, biological processes and molecular processes were identified.

### Principal Component Analysis (PCA)

To check the difference between VSD-placenta and controls, all CpG variables were studied together using principal component analysis. Prior to analysis, we removed all CpG-probes that have missing ß-value followed by remaining ß-value of CpG targets were used for PCA. We used R function “prcomp” to compute principal components (PCs) and used PC1, PC2 and PC3 for the PCA distribution plot. The 3D PCA distribution plot was generated by using R package “ggplot2”.

### Quantitative pyrosequencing

DNA methylation variations were validated on bisulfite-converted genomic DNA by quantitative pyrosequencing. For validation and verification experiments, bisulfite conversion of 1 μg DNA was performed by EZ DNA Methylation GoldTM kit (Zymo Research, Cambridge Bioscience, UK). DNA methylation variations were compared with the data obtained from conventional quantitative pyrosequencing.

The identified differentially-methylated genes were used to generate a heatmap using the Complex Heatmap (v1.6.0) R package (v3.2.2). Ward distance was used for the hierarchical clustering of samples [27].

## Results

Genome-wide DNA methylation analysis of placenta-derived DNA was performed on 8 VSD cases and 10 unaffected controls. Comparison of clinical characteristics between cases and controls are shown in table 1.

### Identification of differentially methylated CpG sites in VSD placenta

We identified 1328 CpG differentially methylated regions in 1328 genes in which there were statistically significant differences (increased or decreased) in cytosine methylation levels in VSD vs control placenta (FDR p-value <0.001). Using the most significantly methylated CpG site as a biomarker for VSD prediction, a total of 80 loci (in 80 genes) had an AUC = 1.00 with significant 95% CI, for VSD detection, Figure 1 (Supplementary Figure 1). A total of 1248 loci (in 1248 genes) had good diagnostic accuracy defined as AUC ≥ 0.81 to 0.99 for VSD detection (Table 1).

**Figure 1.**
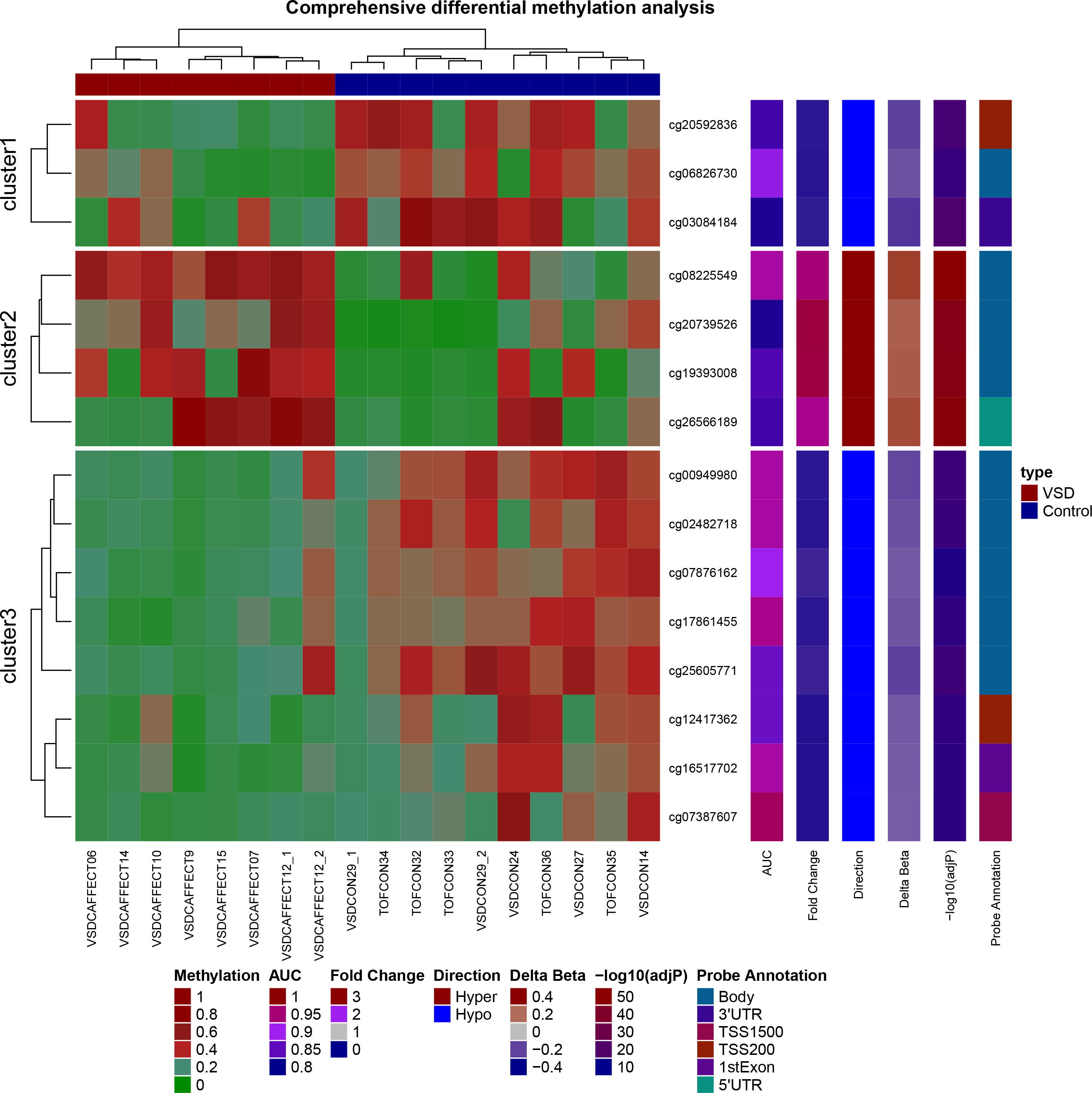
Heatmap of 15 highly differentially methylated loci. Unsupervised hierarchical clustering of differentially methylated loci (rows) in 8 affected and 10 control samples (columns). These 15 CpG loci are at least either 2.0-fold hypomethylated or 2.0-fold hypermethylated with False Detection Rate < 0.00001 in the disease (VSD) condition (red colored squares) compared to normal subjects (blue colored squares). The figure also displays direction, fold change in disease, probe relationship and probe annotation of differentially methylated CpG sites. The red color in the heatmap indicates hyper-DNA-methylation and blue hypo-DNA-methylation values.

**Figure 2.**
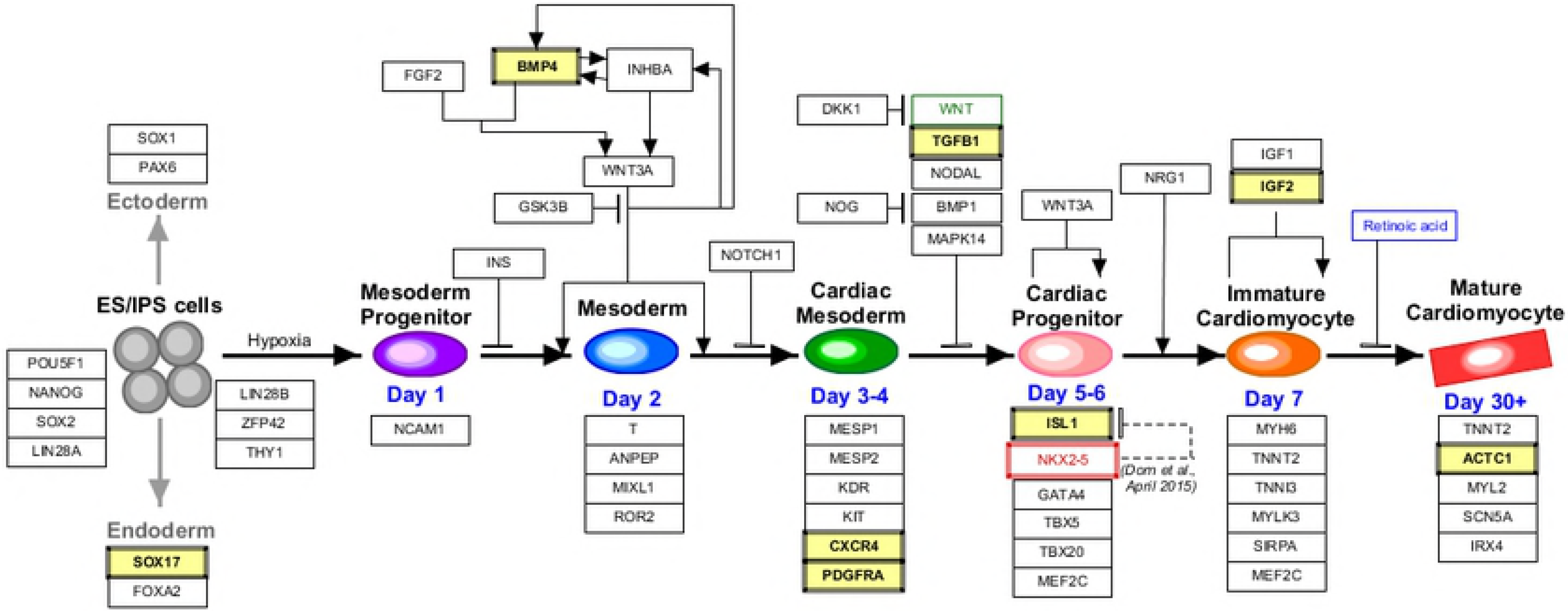
Schematic representation of the cardiac progenitor pathway. Previous studies identified several genetic components related to cardiac development and progression. Genes identified in the present study are shown are highlighted with yellow background.

**Figure 3.**
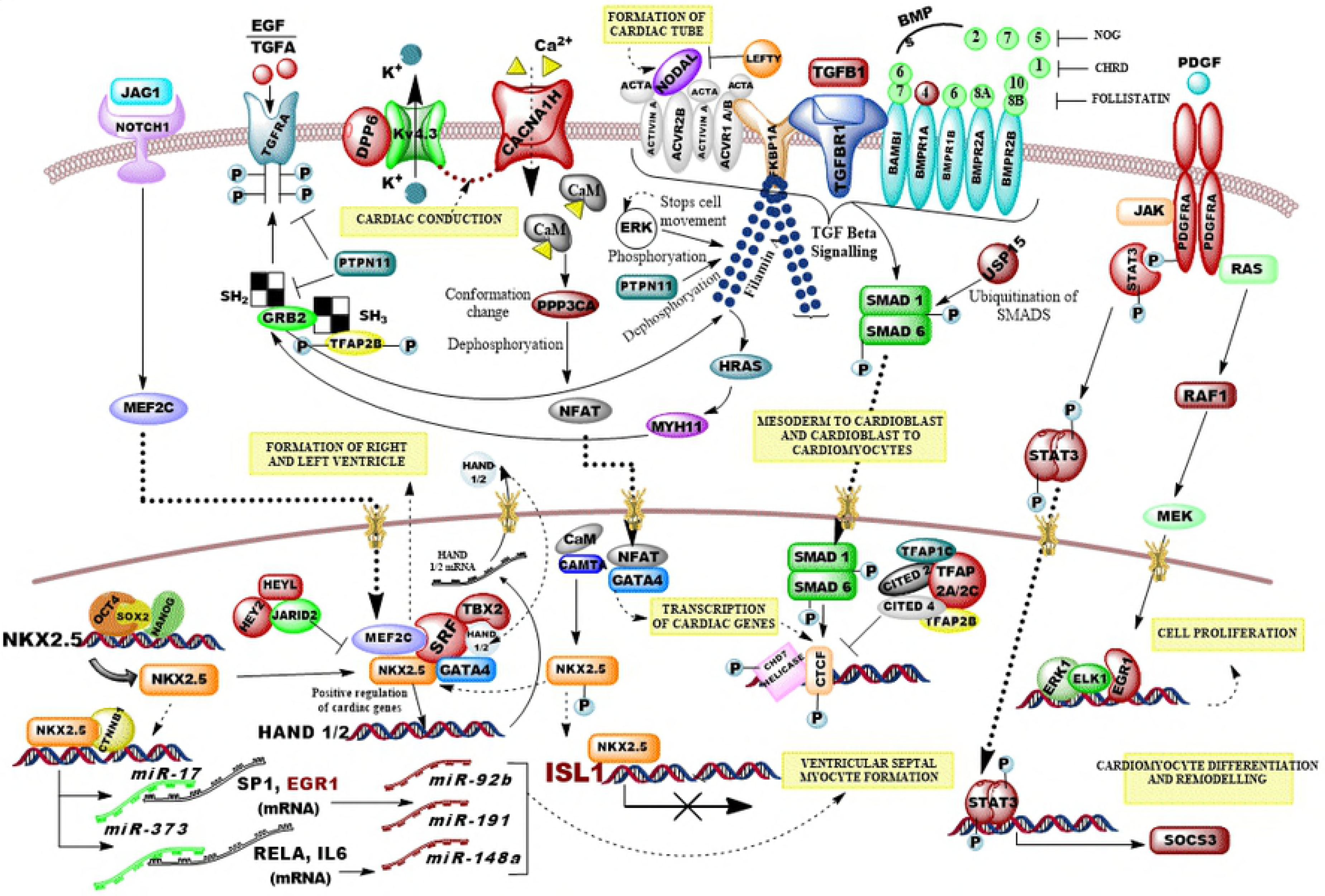
Ingenuity Pathway Analysis (IPA) of significantly differentially methylated genes (at FDR p-value < 0.001). In this study, many important genes were found to be aberrantly methylated in VSD. Various pathways involved in cardiac and ventricular chamber development were over-represented. IPA results indicated the gene network is relevant to cardiac ventricle development, cardiac muscle cell differentiation, embryonic heart tube development, cardiac septum development, heart morphogenesis, cardiac chamber development, and the notch signaling pathway.

Gene ontology analysis revealed an over-representation of pathways known to be involved in cardiac and ventricular chamber development (Figures 2 & 3). Many of the constituent genes in the different pathways were found to have significant dysregulation of DNA methylation. These genes included some known to be responsible for cardiac ventricle (*HEY2*, FDR p=0.002; *ISL1*, FDR p=2.61, E-5), cardiac septum formation (*ISL1* FDR p=2.61, E-5), cardiac muscle development (*ACTC1*: FDR p= 7.15 E-12) and others. Among the 1328 unique genes, the top 10 differentially methylated genes were *BMP4*, *TGFB1*, *PDGFRA*, *TBX2*, *EGR1*, *SRF*, *CXCR4*, *TLX3*, *DPP6*, and *FGFR1*. In addition, the study also identified 8 microRNAs miR-191, miR-548F1, miR-148A, miR-423, miR-92B, miR-611, miR-2110, and miR-548H4 (supplemental Table 1), Significant differential methylation in the genes that coded for 52 Zinc finger proteins (supplemental Table 2), were detected followed by 62 ORFs (open reading frames) (supplemental Table 3) and 28 LOCs (genes without a name and of unknown function) (supplemental Table 4) were detected. Additionally, differential gene methylation for 8 SNOR (Small Nucleolar RNAs) (supplemental Table 5) were identified and two NCRNAs (non-coding RNA) (supplemental Table 6) were also identified. All targets had AUC ROC ≥ 0.81 (FDR p-value <0.001) for the detection of isolated VSD.

Principal Component Analysis (PCA) results displayed a clear-cut difference between VSD and control samples. All VSD samples are away from controls in PCA distribution plot (Supplementary Figure S3) (PCA 3D).

## Discussion

Considering the crucial role of epigenetics in the regulation of gene expression in development, and the increasing evidence linking epigenetic alterations with congenital malformations, we have searched for potential abnormal methylation profiles in placenta DNA of samples with VSD compared to controls. We identified 1488 unique loci, one per gene, that were significantly differentially methylated in the placental DNA in cases of non-syndromic isolated VSD compared to controls. The methylation status of the CpG sites had high predictive accuracy for VSD detection with AUC ≥ 0.80. Indeed 80 of the cytosine loci in 80 genes had AUC = 1.0 and the FDR p-value < 0.005. In addition, significant differential methylation was noted in genes coding for many miRNA, ORFs, NCRAs and LOC’s. These also demonstrated good to excellent diagnostic accuracy for VSD detection. We also identified methylation changes in 8 miRNAs that modulate the activity of 160 genes identified. Pathway analysis identified multiple genes known to be involved in cardiac development overall, ventricular development and cardiac septum formation. These associations give biological plausibility to our findings (Supplementary Table 7).

Cardiac progenitors and their role in heart development: Some of the differentially methylated genes known, suspected or plausibly linked to cardiac development. Recent work indicates that a common multipotent stem cell differentiates into cardiomyocyte, smooth muscle, and endothelial lineages, the major differentiated cell types of the heart [28–30]. This precursor cells are characterized by expression of the transcription factors Isl1 and Nkx2-5. Lineage tracing studies indicate that Isl1 and Nkx2-5 expression contribute to differentiation of the cardiomyocytes and a subset of the smooth muscle and endothelial cells of the heart [30]. Both genes were differentially methylated in CHD cases in our study.

TGF-β signaling in cardiomyocyte differentiation: The TGF-β superfamily members critically regulate many different processes within the cardiovascular system (CVS), including heart development and angiogenesis. The predominant TGF-β superfamily ligands expressed in the CVS are *TGF-β1*, *TGF-β2*, *TGF-β3*, *BMP-2*, *BMP-4*, *BMP-6*, and BMP-7 [31]. The importance of the *TGF-β* superfamily in cardiovascular development is demonstrated by the significant phenotypic changes in knock-out mice models. Disturbances in TGF-β signaling in mice causes embryonic lethality and develop VSDs, myocardial thinning, and a double outlet right ventricle (DORV) a cardiac anomaly in which VSD is a feature [32]. Moreover, BMP-dependent activation of transcription factors including the cardiac zinc finger transcription factor *GATA4*, homeobox protein *NKX2.5*, and myocyte enhancer factor 2C (MEF-2C) augment differentiation mediated by the SMAD signaling pathways that regulate cardiomyocyte proliferation [33].

MicroRNAs and Cardiac diseases: MicroRNAs (miRNAs) are endogenous, small, evolutionary conserved, non-coding RNAs molecules about 22 nucleotides in length that function in the regulation of gene expression [34]. They bind to and inactivate mRNA thus inhibiting protein production. miRNAs are commonly found in clusters through many different regions of the genome, most frequently within intergenic regions and the introns of protein-coding genes [35]. Altered miRNAs expression is known to occur in various cardiovascular disorders such as hypertension, congestive heart failure, CHD, coronary artery disease and stroke. In the present study, significant changes in the methylation of several miRNA genes: miR-191, miR-548f1, miR-148a, miR-423, miR-92b, miR-611, miR-2110, and miR-548h4 gene methylation were observed in the placenta of VSD subjects as compared to controls. Our findings are supported by previous publications indicating these microRNAs are associated with cardiac disorders including VSD and heart failure [36–38].

Chen et al [39] suggested that miR-92b is involved in visceral and cardiac muscle differentiation and regeneration. Additionally, significant changes in miR-92b expression levels have been reported in neointima formation in a rat model of vascular injury [40], and in individuals with heart failure [41], and during cardiac rehabilitation following surgical coronary revascularization [42], suggesting its involvement in these processes. In the present study, a significant variation in placental gene methylation was observed in VSD cases compared to controls.

Fetal / Embryonic cardiac muscle formation: *ACTC1* (Actin, Alpha, Cardiac Muscle 1) gene on chromosome 15q14 (MIM 102540) is the only actin expressed in embryonic heart muscle [43]. It has been suggested that the lack of *ACTC1* may induce apoptosis leading to disrupted cardiac differentiation. Apoptosis plays a crucial role in embryological development and excessive absorption/apoptosis of the primary septum is thought to be a cause of atrioventricular septal defects [44].

Jak / stat signaling in cardiac remodeling and proliferation: One study found that Jak1/Stat3 downstream mediators are induced upon cardiac injury. Inhibition of Stat3 signaling in cardiomyocytes using a dominant-negative transgene, led to fibrotic scarring and reduced cardiomyocyte proliferation [45].

Other significant genes involved in cardiac development: Recent evidence has pointed to *IGF2* as one such molecule. *IGF2* is expressed by epicardial cells during midgestation murine heart development, and IGF2-deficient embryos show reduced cardiomyocyte proliferation in the ventricular wall. Importantly, the cardiomyocyte-specific deletion of the genes encoding the *IGF2* receptors IGF1R and Insr replicated this effect [46].

The inactivation of *Hey2*, a significant Notch signaling molecule, in mice resulted in a spectrum of cardiac malformations that resembled those associated with mutations of human *JAG1*[47] associated with Tetralogy of Fallot (TOF), ventricular septal defects (VSD), and tricuspid atresia. Sakata et al [8] reported that targeted disruption of the *CHF1/Hey2* locus in mice results in VSDs.

Bone Morphogenetic Protein 4 (*BMP4*) on 14q22.2 region. The BMPs are members and the TGF-β signaling family. The effect of BMPs are exerted through receptor-mediated activation of the transcription factor Smad by binding to heterotetrameric receptor complexes [48]. BMPs can induce formation of bone and cartilage, and play a major role in embryogenesis [49]. *BMP4* plays a key role in cardiac development, with expression in ventral splanchnic arteries, branchial-arch mesoderm and outflow tract myocardium underlying the cushion-forming regions of the heart [50, 51]. *BMP4* alterations are known to be associated with defects in cardiac septation such as atrial septal defect (ASD), ventricular septal defect (VSD), and atrioventricular septal defects (AVSD). The inactivation of BMP4 results in neonatal lethality [52, 53].

Isl Lim Homeobox 1 (ISL1) on chromosome 5q11.1. ISL1 is a transcription factor and is important in multipotent cardiovascular cell lineages [54]. The lineages of Isl1-expressing cells include neural crest, endocardium, endothelium, myocardium and smooth muscle cells [55]. Genetic mutations of ISL1 have been found to be associated with CHD and VSD [54, 56].

Platelet-Derived Growth Factor Receptor, Alpha (PDGFR-A). PDGFR-A proteins are receptor tyrosine kinases (RTKs) coded on chromosome 4q12 and is important in PDGF signaling. They specifically localize in the primary cilium of the heart where the downstream effectors P13K-AKT and MEK1/2-ERK1/2 are active and regulate processes like cell cycle control and cell migration in fibroblasts [57, 58]. Cilia play a major role in signaling processes which contribute to the left-right organ asymmetry, differentiation, morphogenesis and the maturation of the heart [58].

*T-BOX 2* (*TBX2*) on chromosome 17q23.2. During heart development, *TBX2* is expressed in the outflow and inflow tracts, inner curvature and atrioventricular canal [59]. Sequence variants at the promoter sequences of *TBX2* have been identified in VSD patients [6].

Early Growth Response 1 (*EGR1*) factor gene on chromosome 5q31.2. *Egr1* controls proliferation, differentiation, and apoptosis of cells, and is also involved in hypoxic and ischemic cardiovascular injuries [60]. *EGR1* is a transcription factor for *NAB1* and is involved in the regulation of physiological and pathological hypertrophy of the heart. *Egr1* may not be required for normal development of muscle mass, but it acquires functional importance during the response to cardiac stress resulting from mechanical overload and hypertrophic signaling [61].

*DPP6* is an essential component of the native cardiac channel complex (Kv4.3, *KCND3*) which regulates cardiac conduction [62]. *DPP6* may serve as an additional beta-subunit responsible for the transient outward current generated during repolarization of the nodal cells in the human heart. Kv4.3, an outward flux potassium ion channel, is expressed extensively in ventricular myocytes.

A significant amount of inactive CaMKII forms molecular complex with Kv4.3 potassium ion channel proteins in cardiomyocytes as a CaMKII reservoir [63]. CaMKII is also a positive regulator of T-Type calcium ion channels, one of which is *CACNA1H* [64]. This channel is responsible for transient calcium ion influx current (I_Ca,T_) into myocardial cells following depolarization[64]. In ventricular myocardium, the T-type Ca^2+^ current channel *CACNA1H*, which is temporarily observed during fetal and neonatal periods, has been shown to reappear in failing/remodeling hearts. These findings suggest that *CACNA1H* is related to cell growth, proliferation, and development [65].

LOC genes. LOC gene(s) are predicted genes whose functions are uncharacterized, and that are currently unnamed. They are assigned to a generic gene and protein category “uncharacterized LOC” plus the GeneID [37]. In the present analysis, we have identified 28 LOC genes on 14 chromosomal regions (supplemental Table 4). Each of these 26 CpG different targets have a ROC AUC ≥ 0.81 and the FDR p-value < 0.005 for VSD prediction. Among these 28 candidates, LOC374443 [66], LOC255512 [67], LOC348926 [68], LOC96610 [69], LOC25845 [70] and LOC220930 [71] expression is associated with different heart ailments. LOC150381 was previously identified under rare Copy Number Variation (CNV) deletion associated with aortic valve stenosis [72].

As previously noted, though DNA methylation patterns are largely tissue specific however recent data suggests correlation between diseases induced methylation changes across tissues [73]. For example, a high degree of correlation in CpG methylation was demonstrated between blood leucocyte and tissue from the lip in cleft lip and palate patients [14]. This phenomenon of cross tissue correlation of methylation markers is the most likely explanation for the observed epigenetic modification in cardiac development genes in the placental of non-syndromic VSD cases.

Our study does have some limitations. One of which is the small sample size. We are now unable to evaluate the role of potential confounders e.g. diet, obesity, and ethnicity on the methylation patterns. Our study however is a proof of concept study whose purpose is to establish the plausibility of a concept. The next step will be to explore this phenomenon in a larger study population.

While it is widely recognized that DNA methylation correlates with gene expression, we did not evaluate this correlation in our study. This is clearly an area for future study. We did however perform GDAC FIREHOSE database analysis which showed that these methylation differences are correlated with altered gene expression in other tissues such as cancer cells (data not shown) (supplemental Table 8, 9) (supplemental Figure 2).

To the author’s knowledge, this is the first study reporting the use of placental molecular markers for CHD detection and for elucidating the mechanism of CHD. We report significant epigenetic dysregulation in the placental DNA of VSD cases. Multiple genes and gene pathways including some known or suspected to be involved in heart and ventricular development were affected. These findings could have scientific and clinical significance in further understanding the pathogenesis of and CHD and in the development of accurate biomarkers using surrogate tissues such as placenta which is accessible through commonly used techniques for analysis. Placental tissue obtained at birth could also be used for the prediction newborn CHD. Newborn screening for critical CHD is now a standard of care.

## Acknowledgments

Contributors: ROB-S conceived and helped design the study, examined the clinical data, supervised the clinical and experimental data interpretation, and critically revised and edited the manuscript. UR analyzed and interpreted molecular and statistical the data and drafted the manuscript. AMV, NKM, CG, performed the statistical and bioinformatics data analysis. SA, RA, RAF, NS participated in drafting the IRB protocol, identified and obtained patient specimens and in reviewing the manuscript. BA, participated in clinical data and specimen collection and reviewing the manuscript. All authors read and critically revised the manuscript and approved the final version.

## Web Resources

The URLs for data presented herein are as follows:

Illumina: http://www.illumina.com/

Genome Studio: http://www.solexa.com/gsp/genomestudio_software.ilmn

Ensemble: http://www.ensembl.org/

UCSC: http://genome.ucsc.edu/

NCBI: http://www.ncbi.nih.gov/

Web Gestalt: http://genereg.ornl.gov/webgestalt/

Chilibot: www.chilibot.net

Ingenuity Systems: www.ingenuity.com).

## Table Headings

**Table 1**. Demographics of VSD cases versus controls

**Table 2**. Differentially methylated genes with Target ID, Gene ID, chromosome location, FDR p-value, and % methylation change in VSD cases and controls for each target methylated. CpG sites with significant FDR p-value indicating methylation status and area under the receiving operator characteristic curve ≥0.80 appear to have strong potential as diagnostic biomarkers for VSD.

## Supplemental Material

**Supplementary Figure S1**. (online only) Receiver operating characteristic (ROC) curve analysis of methylation profiles for four specific markers associated with VSD. The study identified 1328 CpG sites in 1328 genes with significantly differentially-methylated genes that have an area under the ROC curve ≥0.80. At each locus, the False Detection Rate p-value for the methylation difference between VSD subjects and controls was highly significantly different. Due to figure resolution concerns, we have included only four markers (chr 21; cg21161649) (chr 17; cg04245057) (chr 4; cg07809452) (chr 12; cg22129822). AUC: Area Under the Receiver Operating Characteristics Curve; 95% CI: 95% Confidence Interval. Lower and upper confidence intervals are given in parentheses.

**Supplementary Figure S2**. (online only) Open chromatin and transcription factor occupancy in functional CpG sites. Pie charts depicting H3K27Ac histone mark layering, location of CpG sites and transcription factor binding sites.

**Supplementary Figure S2**. (online only). Three dimensional PCA (PCA 3D) of VSD-placenta and control samples. The data are color coded according to their grouping and clustering analysis.

**Supplemental Table S1-S6**. (online only) Differentially methylated CpG sites with Target ID, Gene ID, chromosome location, FDR p-value, and % methylation change in VSD cases and controls for each target methylated for MicroRNA (Supplemental Table S1); Zinc finger protein (Supplemental Table S2); ORF (Supplemental Table S3); LOC (Supplemental Table S4); SNOR (Supplemental Table S5); NCRNAs (Supplemental Table S6).

**Supplementary Table S7**. (online only) Gene Ontology (GO) terms enriched among the genes in the network displayed by GeneMANIA.

**Supplemental Table S8**. (online only) Correlation of Methylation mean with the expression mean in various human tissues from GDAC data. 180 differentially methylated CpG targets were correlated with expression (RNA-seq) data. A bar chart was generated for each CpG target showing the proportion of methylation and mean of expression of the gene in which the CpG target resided.

**Supplemental Table S9**. (online only) Open chromatin conformation and transcription factor binding in differentially methylated CpG sites indicated their role in transcription initiation. ENCODE data showing the H3K27Ac layering on each CpG site presenting an open chromatin conformation. These CpG targets were also occupied with various transcription initiation factors, mostly PolR2A. The position of each CpG site was also noted in respect to the gene in which it resided. Some of the differentially methylated CpG sites that were resided in intronic or 1^st^ exonic regions, signifying their essential function in modulating transcription.

